## Supplementary material for "Robotic platform for microinjection into single cells in intact tissue"

### SUPPLEMENTARY FIGURES AND NOTES

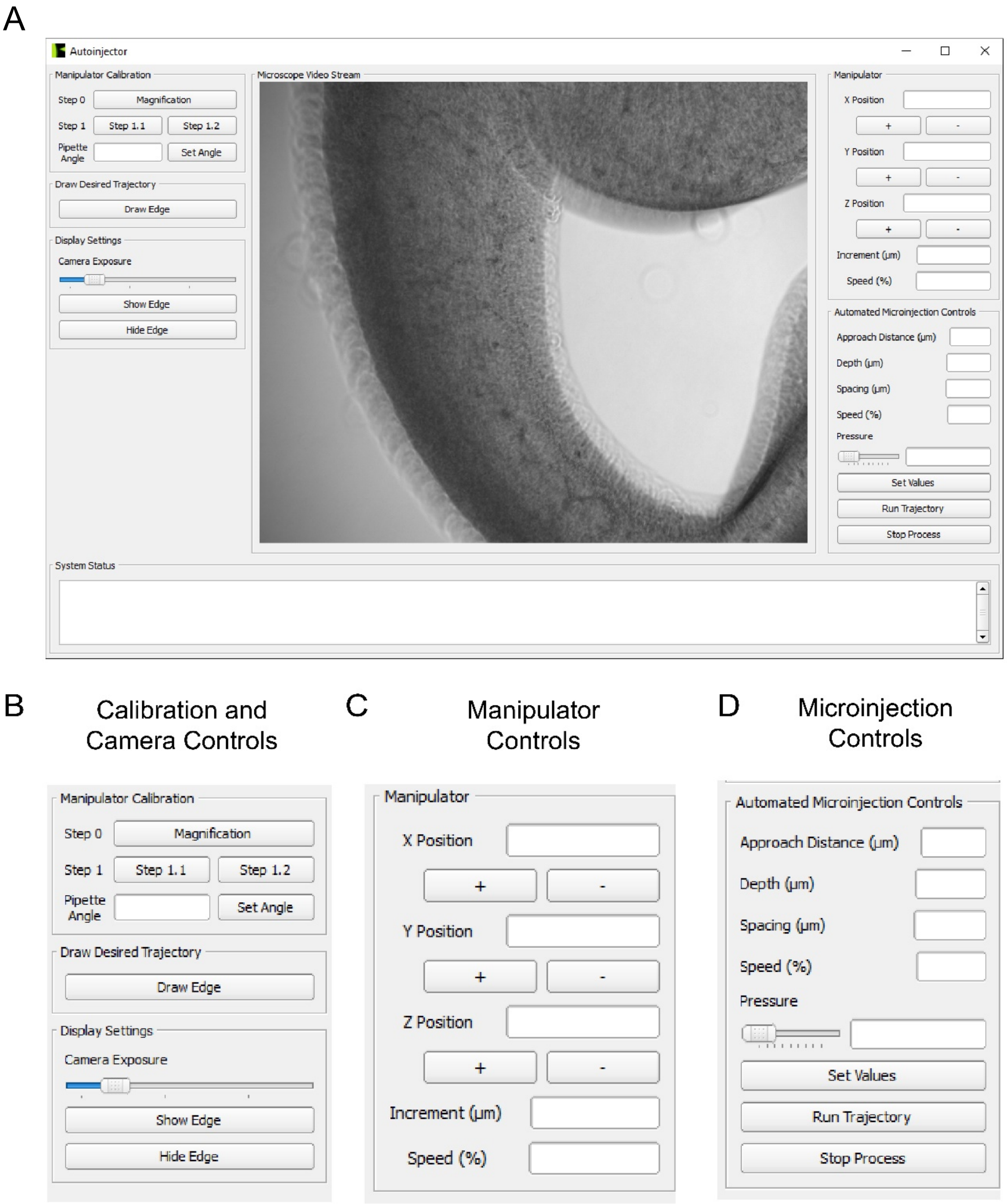

**Supplementary Figure 1: Autoinjector software graphical user interface (GUI).** The GUI developed in Python controls the Autoinjector hardware and allows the user to customize the microinjection protocol. (A) Screenshot of the GUI. Images acquired by the microscope camera are displayed in real time at the center of the GUI. The controls of the microinjection are on the left and right columns of the image. The status of the Autoinjector is displayed in the bottom of the GUI in the response monitor. (B) Controls on the left side of GUI. ‘Calibration Controls’ are used by the user to enable software control of the manipulator in 3D space using the 2D microscope camera images (see **Supplementary Figure 2A** and **Supplementary Note 1** for additional details). The ‘Manual Feature Selection’ controls allows the user to annotate the microscope camera images to define the desired path of microinjection. ‘Single Injection Controls’ allow the user to perform a single injection at a given pressure with a given pulse duration. The ‘Video Capture’ controls allow the user to record the real time microscope camera images as a video. The ‘Feature Viewing’ controls allow the user to display or remove the annotated path of microinjection on the microscope camera image feed. (C) Controls on the right side of GUI. ‘Manipulator Controls’ allow the user to see the current position of the manipulator, and to manually advance the manipulator in X, Y, and Z. The ‘Autoinjector Controls’ allow the user to adjust the microinjection parameters such as micropipette compensation pressure, approach distance, depth of injection, spacing between injections, speed of injection, and number of injection attempts (see **Supplementary Figure 2B** and **Supplementary Note 2** for additional details). The ‘Autoinjector Controls’ also allows the user to start the Autoinjector and to stop the process at any moment.

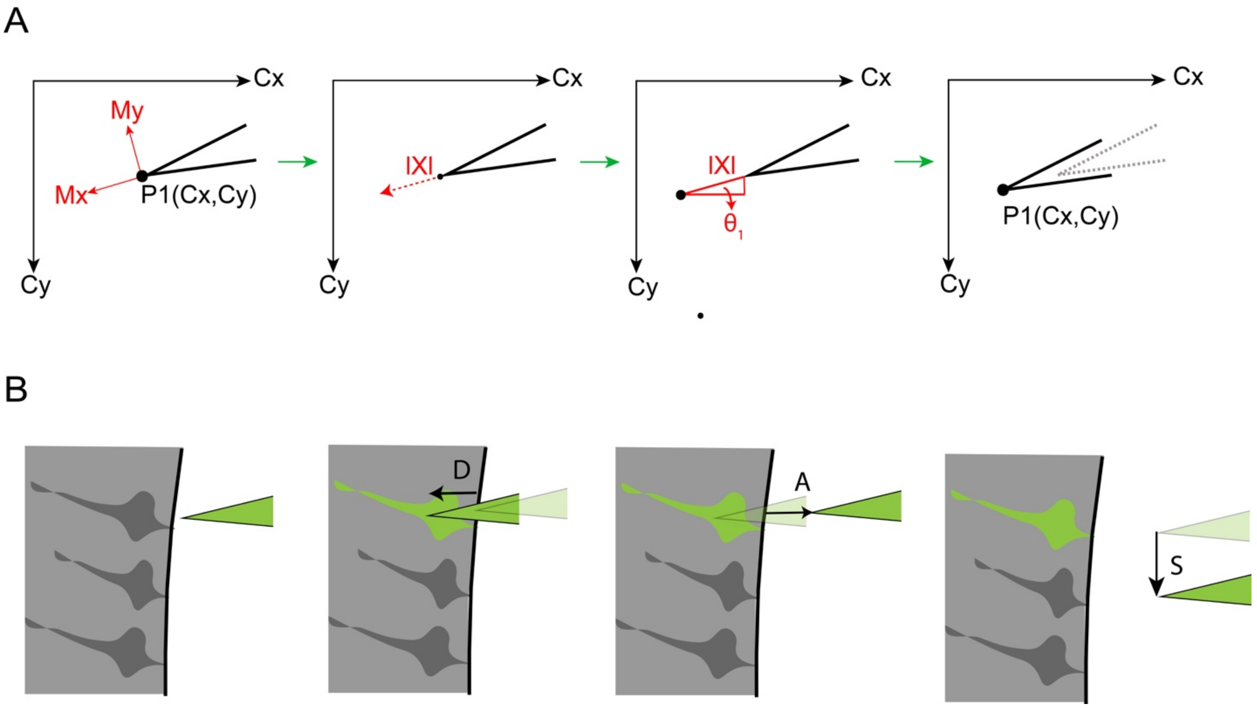

**Supplementary Figure 2: Autoinjector micropipette manipulator calibration and injection parameter definition.** (A) Image guided control of the micropipette required movement in the microscope field of view (FOV) to be translated to the manipulator coordinate system. The micropipette was positioned using the manipulator and had a coordinate system defined by (Mx, My). The tip of the micropipette moved in a second coordinate frame visualized through the microscope FOV defined by (Cx, Cy). The initial position of the micropipette tip in (Cx, Cy) was registered as point P1. The manipulator advanced the micropipette in the manipulator X-axis (Mx) by a predefined distance IXI. The final position of the micropipette tip in (Cx,Cy) was registered as point P2. The distance between P1 and P2 allowed transformation of the displacement IXI to (Cx,Cy) and was used to calculate the angular offset θ between the two coordinate systems. This then allowed programmatically guidance of the micropipette tip to defined points in the camera FOV. (B) The parameters defined by the user to create the final trajectory of the Autoinjector are shown. The micropipette entered a depth, denoted D, into the tissue to perform the microinjection. Next, the micropipette was pulled out a distance termed the approach distance, denoted A. The micropipette was then moved to the next site of injection by moving along the tissue by a distance termed spacing, denoted S. This process was repeated N times where N was the number of injection attempts.

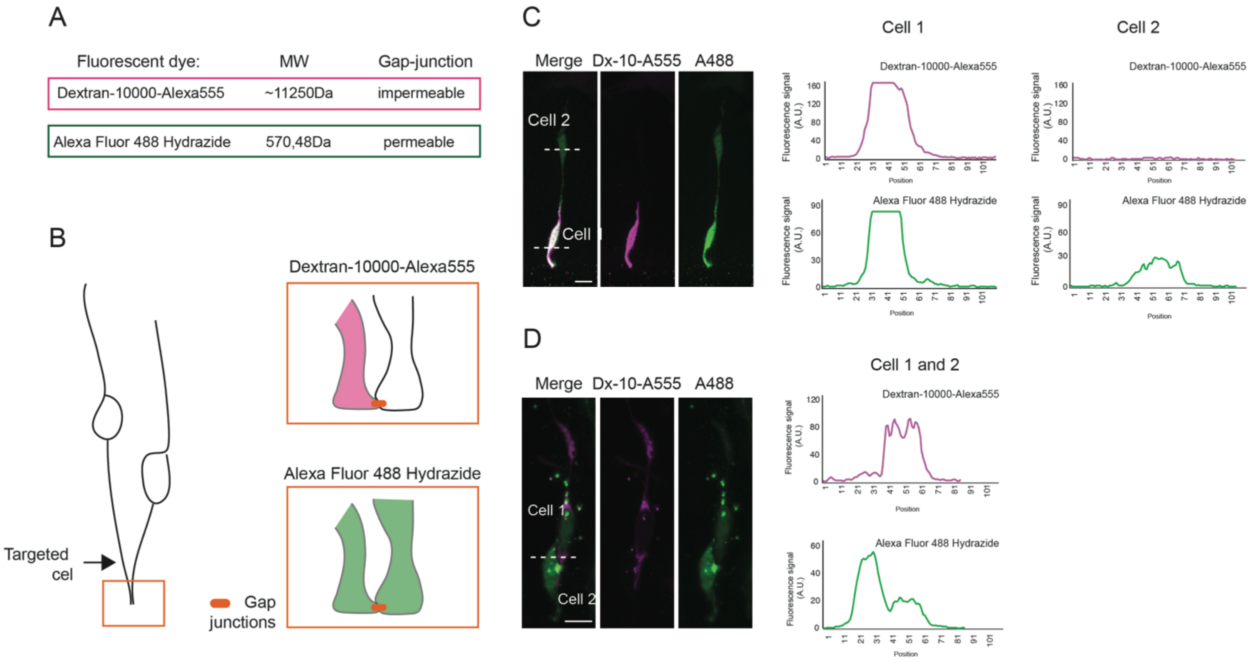

**Supplementary Figure 3: Use of different fluorescent dyes to visualize gap-junction-coupled clusters.** Fluorescent dyes with different molecular weight used to detect gap-junctions-coupled clusters. (A) Molecular weight (MW) and gap-junction permeability of the two fluorescent dyes Dx-10-Alexa555 and Alexa-488. (B) Representative scheme of a 2-cell cluster (cartoon on left). The injected cell is positive for both the junctional impermeable dye Dx-10-Alexa555 and for the junctional permeable dye Alexa-488 (cartoon on the top right, apical endfoot, magenta), while the coupled cell is positive for the junctional permeable dye Alexa-488 only (cartoon on the bottom right, apical endfoot, green). (C, D) Images from 2-cell clusters (left) with the respective line plot quantifications of the fluorescence intensity of Dx-10-Alexa555 and Alexa-488 (right). The line plot was executed along the white dotted lines. Note that the cells shown in C are the same shown in Figure 5B. Scale bars, 10µm.

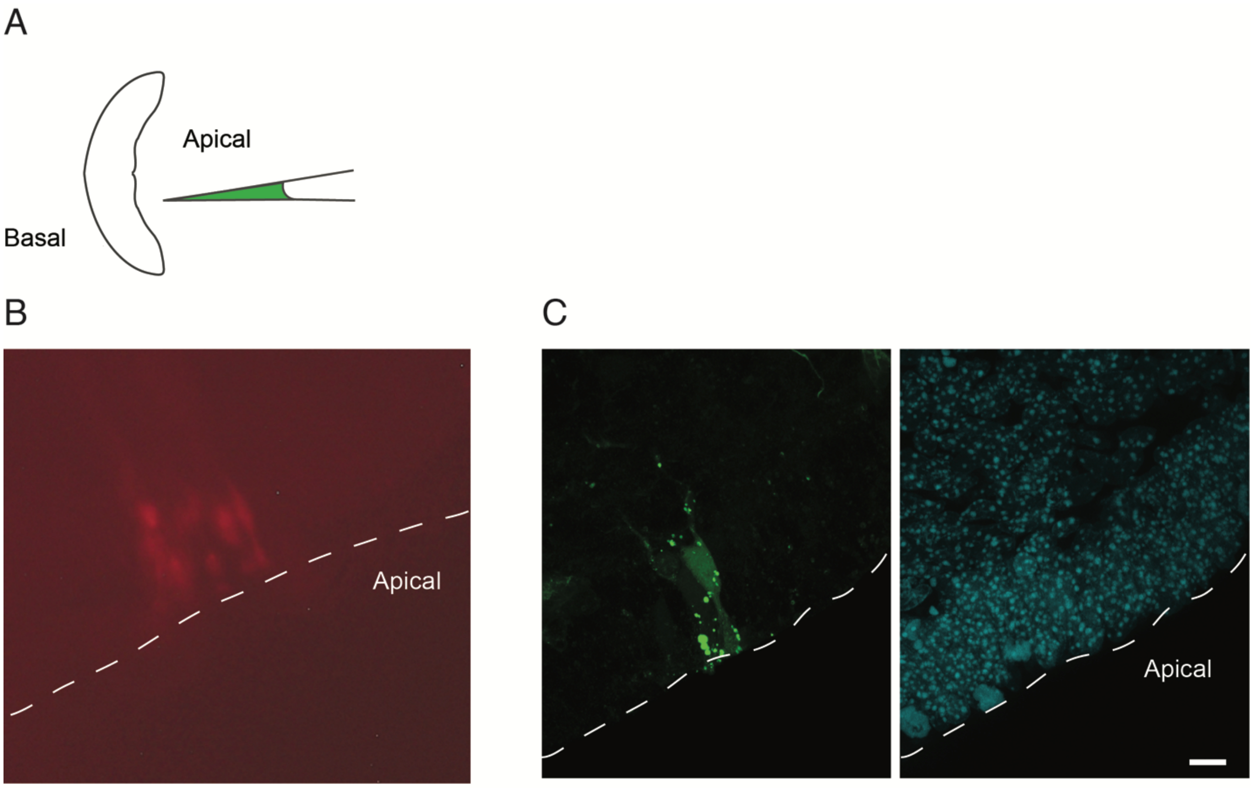

**Supplementary Figure 4: Automated injection of mouse hindbrain APs.** Automated microinjection was performed on organotypic slices of mouse E14.5 hindbrain using Dextran-A555 (red, B) or Dextran-A488 (green, C), followed by fixation (C). (A) Schematic of hindbrain autoinjection. (B) Phase contrast picture of a hindbrain slice immediately after injection. (C) Injected slice fixed and stained for microinjection dye (green), and DAPI (cyan). In both B and C, the injected cells show the typical bipolar morphology of APs. White dotted line: apical surface. Scale bar: 10µm.

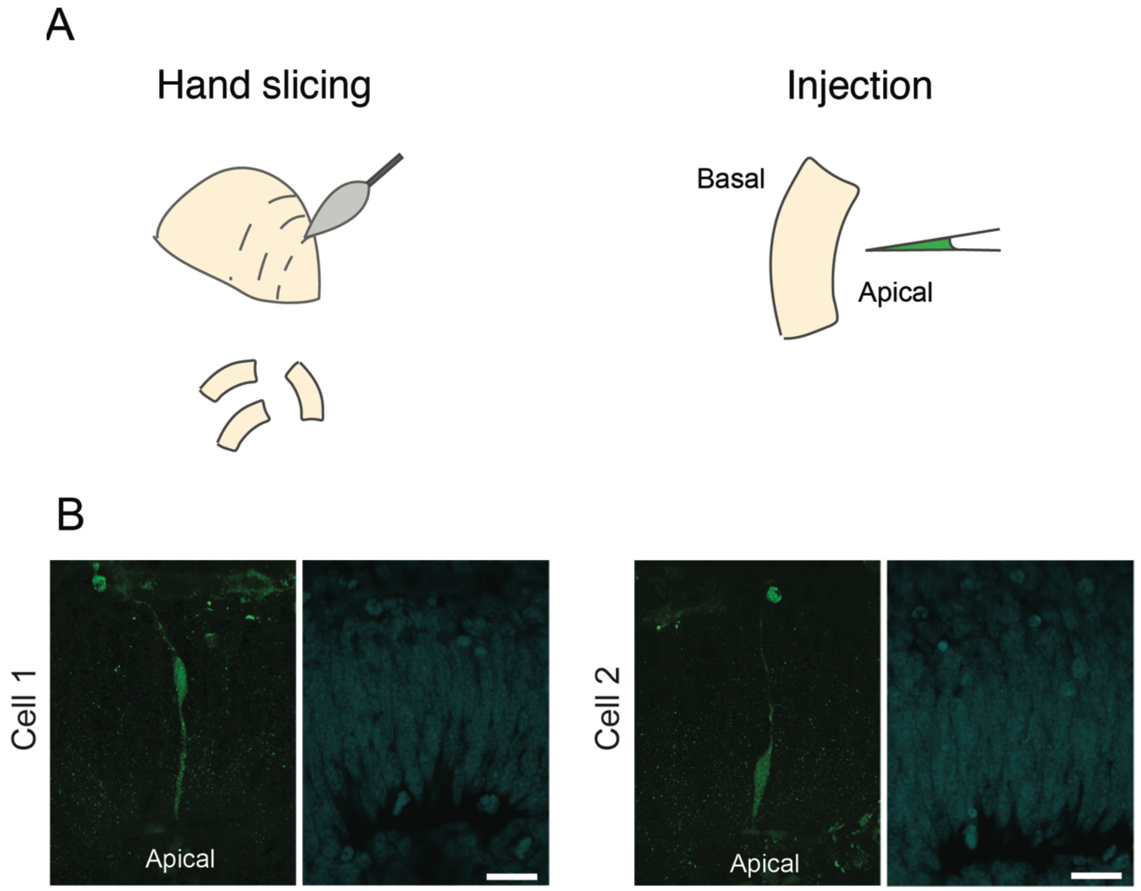

**Supplementary Figure 5: Automated injection of human brain APs.** Automated microinjection was performed on organotypic slices of 12wpc (weeks post conception) human developing telencephalon (A, B) using Dextran-A488 (green, B), followed by fixation and DAPI staining (blue, B). (A) schematic of human tissue hand slicing (top) and auto injection (bottom). (B) two representative examples of injected APs in the human developing telencephalon. The injected cells show the typical bipolar morphology of APs. Scale bars: 20μm.

**Supplementary Note 1: Calibration Procedure**

Movement of the micropipette using the manipulator resulted in displacement of the micropipette in the microscope camera FOV. To control the micropipette to specific locations in the microscope camera field of view (FOV), it was necessary to translate movement of the manipulator in 3D cartesian coordinate space to corresponding change in position of the micropipette in the camera FOV. This translation allowed us to guide the micropipette to any point in the FOV for targeted microinjection. To this end, we performed a calibration procedure at the beginning of each experiment.

A calibration function translated points in the camera axes (Cx, Cy) to points in the manipulator axes (Mx,My) (see **Supplementary Fig. 2A** for a representation of the calibration procedure).

We first needed to determine the angle offset between the manipulator axes and the camera axes.

This was accomplished as follows:

First, the location of the micropipette tip in the camera FOV was recorded and denoted as:

$$P_1 = (C_x, C_y)$$

The manipulator was then advanced along the manipulator X-axis ($M_x$) by a predefined distance $|X|$, resulting in a final position of the micropipette tip in the camera FOV, recorded and denoted as:

$$P_2 = (C_x, C_y)$$

The distance in pixels between $P_1$ and $P_2$ was given by:

$$D(C_x, C_y) = \left|P_2(C_x, C_y) \text{-} P_1(C_x, C_y)\right|$$

The angle between the manipulator X-axis (Mx) and the camera X-axis Cx was given by the equation:

$$\theta = \tan^{-1} \frac{D(C_x)}{D(C_y)}$$

The absolute distance (D) between P1 and P2 was then calculated using the equation:

$$\mathrm{D} = \sqrt{D(C_x)^2 + D(C_y)^2}$$

The scaling factor (S) between manipulator distance and camera pixels was calculated as S:

$$\mathrm{S} = \frac{\mathrm{D}}{|X|}$$

The equation to relate the camera axes to manipulator axes was given by:

$$\begin{bmatrix} C_x \\ C_y \end{bmatrix} = R(\theta) \begin{bmatrix} M_x \\ M_y \end{bmatrix}$$

Where:

$$R(\theta) = \begin{bmatrix} -S\cos\theta & S\sin\theta \\ -S\sin\theta & -S\cos\theta \end{bmatrix}$$

Solution for the manipulator axes in terms of the camera axes was obtained using the equations:

$$\begin{bmatrix} M_x \\ M_y \end{bmatrix} = R^{-1}(\theta) \begin{bmatrix} C_x \\ C_y \end{bmatrix}$$

Where

$$R^{-1}(\theta) = \frac{1}{S^2} \begin{bmatrix} -S\cos\theta & -S\sin\theta \\ S\sin\theta & -S\cos\theta \end{bmatrix}$$

Thus, $M_x$ and $M_y$ were given by:

$$M_x = \frac{1}{S}\left(-C_x\cos\theta - C_y\sin\theta\right)$$

And

$$M_y = \frac{1}{S}\left(C_x\sin\theta - C_y\cos\theta\right)$$

Using these equations, any point in the camera axes ($C_x$, $C_y$) could be converted to a point in the manipulator axes ($M_x$,$M_y$) for image guided position control of the injection micropipette.

**Supplementary Note 2: Trajectory Generation**

The Autoinjector software allowed the user to define a path of microinjection, and to customize the trajectory of microinjection using user defined parameters (see **Supplementary Fig. 2B**). These include: the depth the micropipette is inserted into tissue during microinjection, denoted as D, the distance the micropipette pulls out of the tissue after microinjection attempts, denoted as A, the spacing along the path between subsequent microinjection attempts, denoted as S, the speed the micropipette is inserted into the tissue during the whole procedure, defined as percentage of the maximum speed the manipulator was capable of performing at (1 mm/s), These parameters were utilized to generate the path for the micropipette to take during microinjection was created according to the following protocol:

The user traced a path using the cursor on the camera image along the desired path of microinjection, during which a set of points at different pixels along this path were recorded, This set of pointed was denoted as $t(C_x, C_y)$. A univariate spline interpolation [1,2] was used to connect these points into a continuous path. The interpolated path was denoted as $T(C_x, C_y)$.

Using the calibration procedure (see **Supplementary Note 1**), $T(C_x, C_y)$ was transformed to a trajectory in the manipulator coordinate space. This was denoted as $T(M_x, M_y)$:

$$\begin{bmatrix} T(M_x) \\ T(M_y) \end{bmatrix} = R^{-1}(\theta) \begin{bmatrix} T(C_x) \\ T(C_y) \end{bmatrix}$$

Thus, $T(M_x)$ and $T(M_{y)}$ were given by:

$$T(M_x) = \frac{1}{S}\left(-T(C_x)\cos\theta - T(C_y)\sin\theta\right)$$

and

$$T(M_y) = \frac{1}{S}\left(T(C_x)\sin\theta \text{-} T(C_y)\cos\theta\right)$$

Where $R(\theta)$ was the calibration matrix described in **Supplementary Note 1.**

To compute the final trajectory of microinjection, the coordinates for all injection sites was denoted as:

$$T_i(M_x, M_y) = \begin{bmatrix} T_1(M_x, M_y) \\ T_2(M_x, M_y) \\ \cdots \\ \cdots \\ T_N(M_x, M_y) \end{bmatrix}$$

where and N is the total number of injection attempts. For each point i, the injection micropipette first went to the position $T_i(M_x, M_y)$. The injection micropipette was then moved to $T_i(M_x + D, M_y)$ where D was the user specified the depth of injection. The injection pipette was then moved to $T_i(M_x + D\text{-}A, M_y)$, resulting in a retraction of the micropipette by a distance A. The micropipette was then moved to $T_i(M_x + D, M_y\text{-}s)$, where s was the user defined inter injection site spacing. This process was repeated until the end of the drawn line was reached.

**Supplementary Note 3: Calibration consistency across users**

Visual detection of pipette tip during the calibration can depend on individual perception and can subsequently be a source of positioning error of the Autoinjector. We examined user variability by having four independent users perform the calibration which solves for the angle offset between the manipulator axes and the camera axes. One of the users was an expert user of the Autoinjector, and the other three users had no prior experience calibrating the device. The angle offset calculated from the expert user's calibration was found to be 2.99 ± 0.163 degrees (n = 3 calibrations). The angle offset values calculated from the inexperienced user's calibrations were found to be 2.64 ± 0.223 degrees for inexperienced user 1, 2.86 ± 0.382 degrees for inexperienced user 2, and 2.73 ± 0.341 degrees for inexperienced user 3. Significance tests were performed using a Welch's t-test indicated that there was no significant difference between the experienced user and inexperienced user 1 (p = 0.2817), inexperienced user 2 (p = 0.5089), nor inexperienced user 3 (p = 0.5311). To determine the effects of user calibration on resolution, we performed a resolution test described in the main text (except for here using 4 points rather than 8) using the inexperienced user's calibration angle with the highest standard deviation (inexperienced user 2, calibration angle = 2.86 ± 0.382 degrees). The inexperienced user 2 had an average error in the manipulator axes for the x, and y axis of 0.524 ± 0.409 µm, and 0.518 ± 0.480 µm, respectively (n = 5 measurements per point, 4 points). This value was deemed sufficient for experiments as the distances used in experiments are much greater than this resolution value. Thus, the accuracy obtained across experimenters indicated that variation in user's visual perception does not significantly affect calibration angles.

**Supplementary Table 1: Parts and Reagents**

| REAGENT or RESOURCE | SOURCE | IDENTIFIER |
|---|---|---|
| **Antibodies** | | |
| Chicken anti-Tbr2 | Merck Millipore | Cat# AB15904, RRID: AB_10615604 |
| Rabbit anti-Tbr2 | Abcam | Cat# ab222226, RRID: AB_2721040 |
| Rabbit anti-Tbr2 | Abcam | Cat# ab183991, RRID: AB_2721040 |
| Rabbit anti-RFP | Rockland antibodies | Cat# 600-401-379, RRID: AB_2209751 |
| Rabbit anti-g-tubulin | Sigma-Aldrich | Cat# T5192, RRID: AB_261690 |
| Mouse anti-g-tubulin | Sigma-Aldrich | Cat# T6557, RRID: AB_477584 |
| Mouse anti-TUJ1 | BioLegend | Cat# 801201, RRID: AB_2313773 |
| Mouse anti-TUJ1, Alexa Fluor®488 | BioLegend | Cat# 801203, RRID: AB_2564757 |
| Mouse anti-TUJ1, Alexa Fluor®647 | BioLegend | Cat# 801209, RRID: AB_2686930 |
| Mouse anti-Dextran | STEMCELL Technologies | Cat# 60026, RRID: AB_2651016 |
| Mouse Anti-Dextran, FITC | STEMCELL Technologies | Cat# 60026FI, RRID: AB_2651016 |
| Mouse anti-Reelin | Abcam | Cat# ab78541, RRID: AB_1603148 |
| Lucifer Yellow Polyclonal Antibody | Thermo Fisher Scientific | Cat# A-5750, RRID: AB_2536190 |
| Alexa Fluor 488Polyclonal Antibody | Thermo Fisher | Cat# A-11094, RRID: AB_221544 |
| Goat anti-Chicken IgY (H+L) Secondary Antibody, Alexa Fluor 647 | Thermo Fisher Scientific | Cat# A-21449, RRID:AB_2535866 |
| Donkey anti-Mouse IgG (H+L) Secondary Antibody, Alexa Fluor A555 | Thermo Fisher Scientific | Cat# A-31570, RRID: AB_2536180 |
| Donkey anti Mouse IgG (H+L) Secondary Antibody, Alexa Fluor A488 | Thermo Fisher Scientific | Cat# A-21202, RRID: AB_141607 |
| Donkey anti Mouse IgG (H+L) Secondary Antibody Alexa Fluor A647 | Thermo Fisher Scientific | Cat# A-31571, RRID: AB_162542 |
| Donkey anti Rabbit IgG (H+L) Secondary Antibody, Alexa Fluor A488 | Thermo Fisher Scientific | Cat# A-21206, RRID: AB_2535792 |
| Donkey anti Rabbit IgG (H+L) Secondary Antibody, Alexa Fluor A647 | Thermo Fisher Scientific | Cat# A-31573, RRID: AB_2536183 |
| Donkey anti Rabbit IgG (H+L) Secondary Antibody Alexa Fluor A555 | Thermo Fisher Scientific | Cat# A-31572, RRID: AB_162543 |

| | | |
|---|---|---|
| **Bacterial and Virus Strains** | | |
| One Shot TOP10 Chemically Competent E.coli | Thermo Fisher Scientific | Cat# C404010 |
| **Biological Samples** | | |
| 12 wpc human fetal neocortical tissue | Tissue obtained with informed written maternal consent, followed by elective pregnancy termination from Klinik und Poliklinik für Frauenheilkunde und Geburtshilfe, Universitätsklinikum Carl Gustav Carus of the Technische Universität Dresden | N/A |
| **Chemicals and Peptides** | | |
| DAPI (4′,6-Diamidine-2′-phenylindole dihydrochloride) | Thermo Fisher Scientific | Cat# 000000010236276001 |
| Alexa Fluor 647 Phalloidin | Thermo Fisher Scientific | Cat# A22287 |
| Alexa Fluor 488 Phalloidin | Thermo Fisher Scientific | Cat# A12379 |
| Alexa Fluor 488 Hydrazide | Thermo Fisher Scientific | Cat# A10436 |
| Dextran, Alexa Fluor 488; 3,000 MW, Anionic | Thermo Fisher Scientific | Cat# D34682 |
| Dextran, Alexa Fluor 488; 10,000 MW, Anionic | Thermo Fisher Scientific | Cat# D22910 |
| Dextran, Alexa Fluor 555; 10,000 MW, Anionic | Thermo Fisher Scientific | Cat# D34679 |
| Lucifer Yellow CH, Lithium Salt, for microinjection | Cat# L12926 | |
| mMessage mMachine T7 Ultra Transcription Kit | Cat# AM1345 | |
| Agarose, Low Melt | Carl Roth | Cat# 6351.2 |
| Agarose, Wild Range | Sigma-Aldrich | Cat# A2790 |
| B-27 Supplement | Thermo Fisher Scientific | Cat# 17504044 |
| Cellmatrix Type-IA (Collagen, Type !) | FUJIFILM Wako Chemicals | Cat# 637-00653 |
| Distilled Water | | |
| DMEM-F12, CO2 independent (w/o Phenol red) | Sigma-Aldrich | Cat# D2906 |
| DMEM-F12, CO2 independent (with Phenol red) | Sigma-Aldrich | Cat# D8900 |
| HEPES-NAOH, pH 7.2, 1M (HEPES buffer) | Carl Roth | Cat# 9105.3 |
| L-Glutamine, 200 mM | Thermo Fisher Sientific | Cat# 25030024 |
| Liquid Nitrogen | AirLiquide | |
| Mowiol 4-88 | Sigma-Aldrich | Cat# 81381 |
| N-2 Supplement | Thermo Fisher Scientific | Cat# 17502048 |
| Neurobasal Medium | Thermo Fisher Scientific | Cat# 21103049 |
| Nuclease-free water | Thermo Fisher Scientific | Cat# AM9937 |
| Rat serum | Charles River Laboratories Japan | |
| Best-CA 221 Glue | Best Klebstoffe GmbH & Co.KG | Cat# CA221-10ml |

| | | |
|---|---|---|
| Sodium bicarbonate (NaHCO3) | Merck | Cat# 106323 |
| Sodium hydroxide (NaOH) | Merck | Cat# 106482 |
| Tyrode's salt | Sigma | Cat# T2145-10x1L) |
| Paraformaldehyde | Merck | Cat# 818715 |
| Penicillin-Streptomycin (10,000 U/ml) | Thermo Fisher Scientific | Cat# 15140122 |
| PBS | | |
| O2 (40%), CO2 (5%), N2 (55%) Mix, 50 liters | | |
| **Experimental Models: Organism/Strains** | | |
| Mouse: C57BL/6JOlaHsd | Envigo | N/A |
| Recombinant DNA | Taverna et al., 2001 | |
| pTNT-mRFP | | N/A |
| **Equipment** | | |
| Borosilicate glass capillaries, 1.2 mm outer diameter x 0.94 mm inner diameter | Sutter Instruments | Cat# BF-120-94-10 |
| Bottle-top filter system, 500 ml | Corning | Cat# 430769 |
| Light source | Olympus | Cat# Highlight 3100 |
| Falcon tubes, 15 ml | Corning | Cat# 430791 |
| Falcon tubes, 50 ml | Corning | Cat# 430829 |
| Fine-tip paintbrush | | |
| Flaming/ Brown micropipette puller | Sutter Instruments | Cat# P-97 |
| Forceps, Dumont no. 3 | Fine Science Tools | Cat# 11231-30 |
| Forceps, Dumont no. 5 | Fine Science Tools | Cat# 11255-20 |
| Forceps, Dumont no. 55 | Fine Science Tools | Cat# 11252-20 |
| Microloader Tips | Eppendorf | Cat# 5242956.003 |
| Heating block | Labtech International | Cat # Dri block Digi2 |
| Inverted fluorescence microscope | Zeiss | Cat# Axiovert 200 |
| Motorized stage XY for microscope | | |
| Three-axis Manipulator | Sensapex Inc | Cat# tree-axis uMP |
| Microcontroller | Arduino | Cat# Arduino Due |
| Microscope camera Hamamatsu Orca Flash 4.0 V3 | | |
| Custom pressure rig | | |
| Manual pressure regulator | McMaster Carr | Cat# 0-60 PSI 41795K3 |
| Electronic pressure regulator | Parker Hannifin | Cat# 990-005101-002 |
| Solenoid valve | | Cat# LHDA053321H-A |
| Computer PC | | |
| Pipette holder | | Cat# 64-2354 MP-s12u |
| Multiwell plate, 24 wells | Nunc | Cat# 142475 |
| Pasteur pipettes, plastic | | |
| Pipette and tips | | |

| | | |
|---|---|---|
| Petri dish, 60 x 15 mm | Greiner | Cat# 628102 |
| Petri dish, 35 x 10 mm | Nunc | Cat# 153066 |
| Petri dish, 34 x 14 mm, including Microwell no. 1.5 cover glass | Cat# P35G-1.5-14-C | |
| Puller filament, 3.0-mm square box filament | | Cat# FB330B |
| Slice culture incubation box | MPI-CBG | Cat# custom made |
| Stereomicroscope | | Cat# SZX12 |
| Tabletop centrifuge | | Cat# 5431622 |
| Thermometer | | |
| Vibratome | | Cat# VT1000s |
| Whole-embryo-culture-system incubator | | Cat# RKI-10-0310 |
| Waterbath | | |
| **Software and Algorithms** | | |
| Fiji | | RRID: SCR_002285 |
| Python | Python Software foundation | Python 2.7.12 |
| Arduino | Arduino | |
| ZEN | | RRID: SCR_013672 |
